# A labeled-line for cold from the periphery to the parabrachial nucleus

**DOI:** 10.1101/633115

**Authors:** Junichi Hachisuka, H. Richard Koerber, Sarah E. Ross

## Abstract

Spinal projection neurons are a major pathway through which somatic stimuli are conveyed to the brain. However, the manner in which this information is coded is poorly understood. Here, we report the identification of a modality-selective spinoparabrachial (SPB) neuron subtype with unique properties. Specifically, we find that cold-selective SPB neurons are differentiated by selective afferent input, reduced neuropeptide sensitivity, distinct physiological properties, small soma size, and low basal drive. In addition, optogenetic experiments reveal that cold-selective SPB neurons are distinctive with respect to their connectivity, with little to no input from either Pdyn or Nos1 inhibitory interneurons. Together, these data define a neural substrate supporting a labeled-line for cold from the periphery to the brain.

## INTRODUCTION

Cold, along with heat, pain, itch, and some aspects of touch are conveyed from the spinal cord to the brain via the anterolateral tract (Braz et al., 2014; Todd, 2010). This pathway is made up of neurons that arise from many distinct laminae of the spinal cord and project to numerous regions of the brain, ultimately giving rise to autonomic, affective and discriminative aspects of somatosensation. Although numerous groups have recorded from neurons that contribute to the anterolateral tract, the number of distinct spinal output subtypes remains unclear, and how these parallel channels of information give rise to discrete aspects of somatosensation is not known.

To better understand somatosensory coding, it is critical to identify the different channels of output from the spinal cord. In this regard, one of the key challenges is that most spinal output neurons respond to several types of stimuli. For instance, the majority of temperature-responsive spinal output neurons also respond to mechanical input (Allard, 2019; Craig et al., 2001; Craig and Andrew, 2002; Ferrington et al., 1987). Similarly, all spinal output neurons that appear to be tuned for itch also respond to the noxious chemicals mustard oil and capsaicin (Andrew and Craig, 2001; Davidson et al., 2012; Jinks and Carstens, 2002). Finally, wide dynamic range neurons respond to both innocuous and noxious stimuli (Andrew, 2010; Ferrington et al., 1987; Willis et al., 2017). These diverse somatosensory stimuli feel different and engage different autonomic and affective responses, arguing that they must be differentially represented within the nervous system. However, the manner in which this information is encoded by spinal output neurons with polymodal response properties remains a subject of debate.

Here, we focused on one aspect of somatosensation — cold — because there was a strong precedent for the idea this modality is conveyed, at least in part, by spinal output neurons that are modality-selective rather than polymodal. In particular, extracellular recording from cat and monkey spinal cord in vivo has revealed the existence of spinothalamic tract (STT) neurons that respond to cold stimulation to the skin, but not other somatosensory modalities (Craig et al., 2001; Craig and Dostrovsky, 2001; Craig and Hunsley, 1991; Craig and Kniffki, 1985; Dostrovsky and Craig, 1996; Zhang et al., 2006). In rodent, we and others have described spinoparabrachial (SPB) neurons that are cold-selective (Allard, 2019; Andrew, 2009; Hachisuka et al., 2016). In this study, we identified cold-selective output neurons and then further examined whether they show distinctive features indicative of a distinct subtype of SPB neuron. Here, we report that cold-selective SPB neurons are significantly different from other SPB neurons in many regards including neuropeptide responsivity, soma size, physiological character, and microcircuit connectivity. These data suggest that cold is conveyed from the periphery to the parabrachial nucleus via a distinct and specific population of ‘SPB-cold’ neurons.

## RESULTS

### Identification of cold-selective SPB neurons

In order to characterize the response properties of lamina I SPB neurons to natural stimulation to the skin, we used the semi-intact somatosensory preparation (Koerber and Woodbury, 2002; Mcilwrath et al., 2007) that was recently modified for whole-cell recordings of identified spinal cord neurons (Hachisuka et al., 2016). This preparation comprises of a large portion of spinal cord (~C2 – S3), together with L2 and L3 roots, ganglia, saphenous nerve and femoral cutaneous nerve and hind limb skin, dissected in continuum (Figure 1A). Lamina I SPB neurons for whole-cell patch clamp recordings were retrogradely labeled through stereotaxic injections of DiI into the lateral parabrachial nucleus. To functionally characterize these neurons, we measured their action potentials to cold (0 °C saline), moderate mechanical stimulation (firm brush and/or von Frey filaments, 1 – 4 g), and heat (50 °C saline).

**Figure 1.**
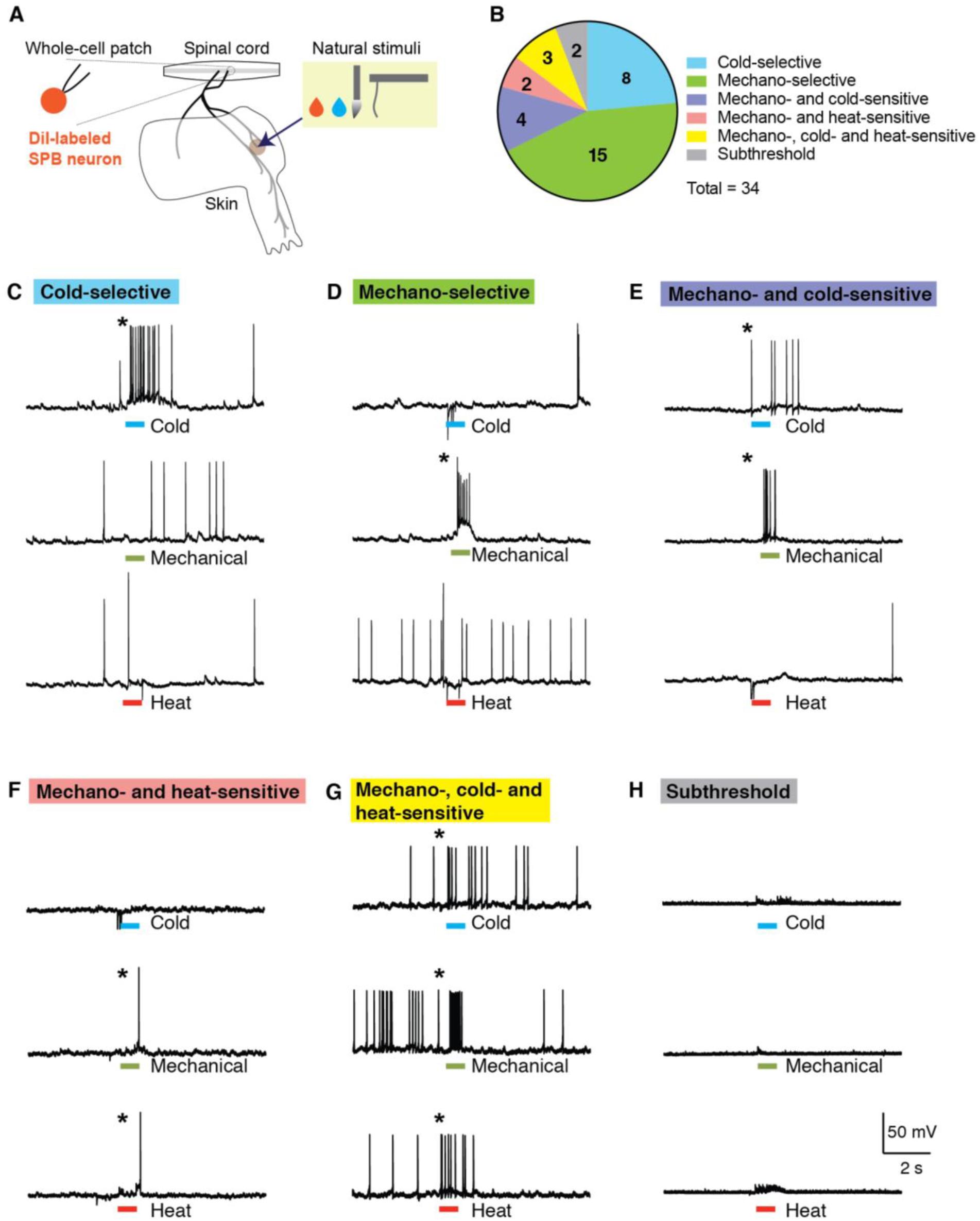
A subset of SPB neurons are cold-selective. **(A)** Schematic of the semi-intact somatosensory preparation. Spinal cord, L2 and L3 roots, saphenous nerve, lateral femoral cutaneous nerve, and hind paw skin are taken together and whole-cell patch clamp recording is made from retrogradely-labeled SPB neurons. (**B)** Pie chart of response properties of SPB neurons to mechanical (small, firm brush and/or von Frey filaments, 1 – 4 g), cold (0°C saline) and heat (50 °C saline) stimulation to the skin. Approximately 25% of SPB neurons are cold-selective. n = 34 SPB neurons. (**C-H)** Example traces of a cold-selective SPB neuron **(C)**, a mechano-selective neuron **(D)** a mechano- and cold-sensitive neuron **(E)**, a mechano- and heat-sensitive neuron **(F)**, a mechano-heat- and cold-sensitive neuron **(G)**, and a subthreshold neuron, which showed EPSPs but no action potentials. * indicates response to stimulation is greater than 3 SD of the baseline activity.

In these initial experiments, we recorded from and characterized 34 retrogradely-labeled lamina I SPB neurons. Consistent with previous studies (Allard, 2019; Andrew, 2009; Bester et al., 2000; Hachisuka et al., 2016), we found numerous mechano-selective neurons (Figures 1B and D). There was also a large subset of neurons that were polymodal, responding to both mechanical and thermal (Figures 1B, 1E, 1F and 1G). None of these SPB neurons responded to heat alone. In contrast, ~25% (8 of 34) responded selectively to cold (Figures 1B and 1C). Finally, two neurons were considered subthreshold because, although they showed evoked excitatory post synaptic potentials (EPSPs) upon stimulation of the skin, this input was not strong enough to evoke an action potential (Figured 1B and 1H).

### Cold-selective SPB neurons receive input from one afferent subtype only: cold afferents

Cold-selective SPB neurons are so-defined by their output—they *fire* in response to cold only. However, whether these cells receive only cold input had never been examined. To examine whether cold-selective SPB neurons receive subthreshold input from other modalities, we recorded evoked EPSCs in response to natural stimulation of the skin. In particular, cold-selective SPB neurons were identified in current clamp mode and then recorded in voltage clamp mode (at the reversal potential of chloride, −70 mV) to permit the isolation of excitatory inputs. As expected, application of cold (0 °C saline) to the skin significantly increased the frequency of excitatory post synaptic currents (EPSCs) onto cold-selective SPB neurons (Figures 2A, 2B and 2C). In contrast, there was no significant increase in the EPSC frequency in response to mechanical stimulation (Figures 2D, 2E and 2F) or heat (Figures 2G, 2H and 2I). These data suggest that cold-selective SPB neurons receive input from only one modality, cold.

**Figure 2.**
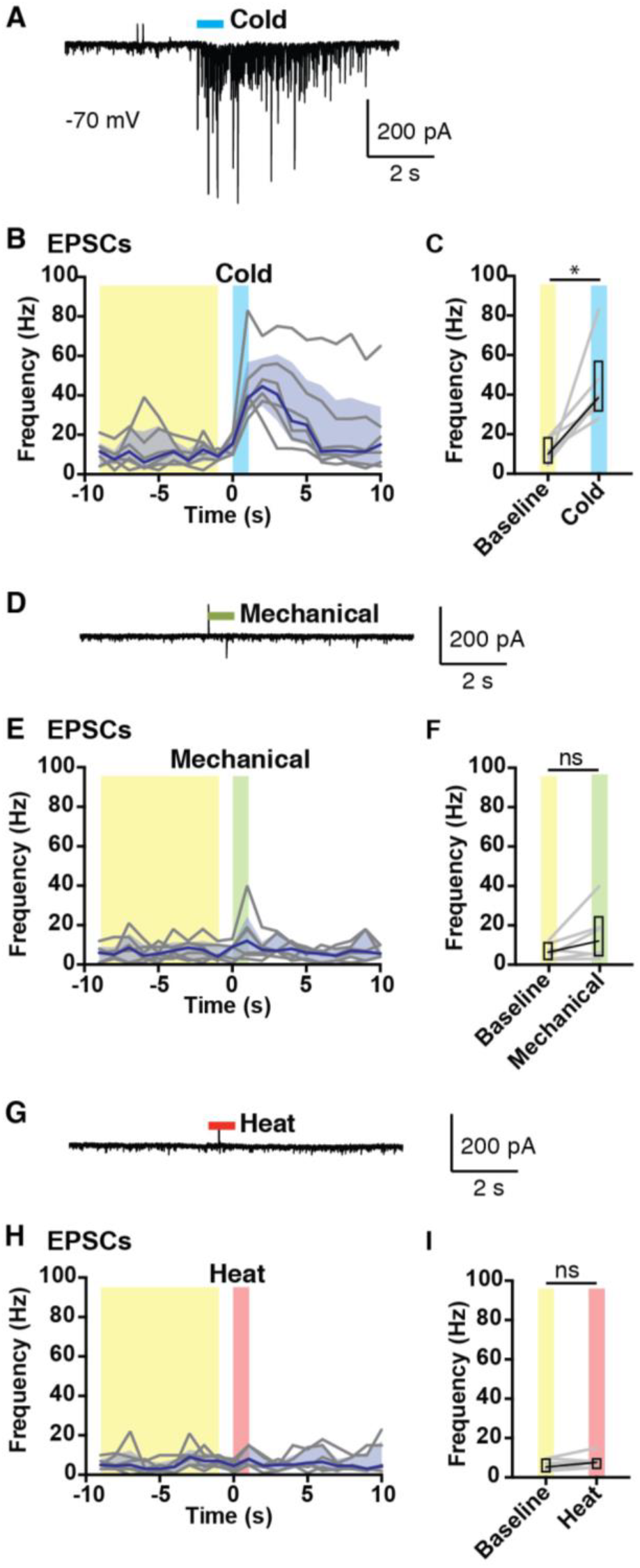
Cold-selective SPB neurons receive only cold input. Analysis of EPSCs of cold-selective neurons evoked by natural stimuli. (**A-C)** Representative trace **(A)** and quantification **(B-C)** of EPSCs observed in response to cold (0° C saline). (**D-F)** Representative trace **(D)** and quantification **(E-F)** of EPSCs observed in response to mechanical stimulation (stiff brush or von Frey filaments, 1, 2 or 4 g)). (**G-I)** Representative trace **(G)** and quantification **(H-I)** of EPSCs observed in response to heat (0° C saline). Black lines are median, shaded area represents interquartile range, and grey lines represent data from individual cells (n = 6 SPB neurons). * indicates EPSC frequency over baseline period (averaged over 9 s, highlighted in yellow) is significantly different than EPSC frequency in response to cold (1 s, highlighted in blue), p < 0.05, Wilcoxon test. For mechanical (highlighted in green) and heat (highlighted in red), n.s. indicates not significant, p > 0.05, Wilcoxon test.

### Cold-selective neurons are unresponsive or only weakly responsive to Substance P

It has previously been shown that the majority of SPB neurons express Tacr1 (also known as the Neurokinin 1 Receptor), which is the receptor for Substance P (also known as Tac1) (Cameron et al., 2015). We therefore assessed whether cold-selective SPB neurons contained functional Tacr1 to determine whether they belong to this category. Sixteen retrogradely-labeled SPB neurons were studied in voltage clamp mode, and neurons were considered to express functional Tacr1 if an inward current was induced by bath-applied Substance P (2 µM). Interestingly, cold-selective SPB neurons showed little to no inward current in response to Substance P (Figures 3A and 3C). In contrast, the majority of other lamina I SBP neurons (i.e., those that were not cold-selective) showed strong inward current to Substance P (Figures 3B and 3C). Overall, the response of cold-selective SPB neurons was significantly smaller than that of other SPB neurons (Figures 3B and 3C), suggesting that cold-selective SPB neurons express little to no functional Tacr1. Since the majority of noxious information is thought to be conveyed by Tacr1-expressing SPB neurons (Doyle and Hunt, 1999; Khasabov et al., 2002; Labrakakis and MacDermott, 2003; Mantyh et al., 1997), this finding supports the idea that nociceptive output and cold output are mediated via distinct channels.

**Figure 3.**
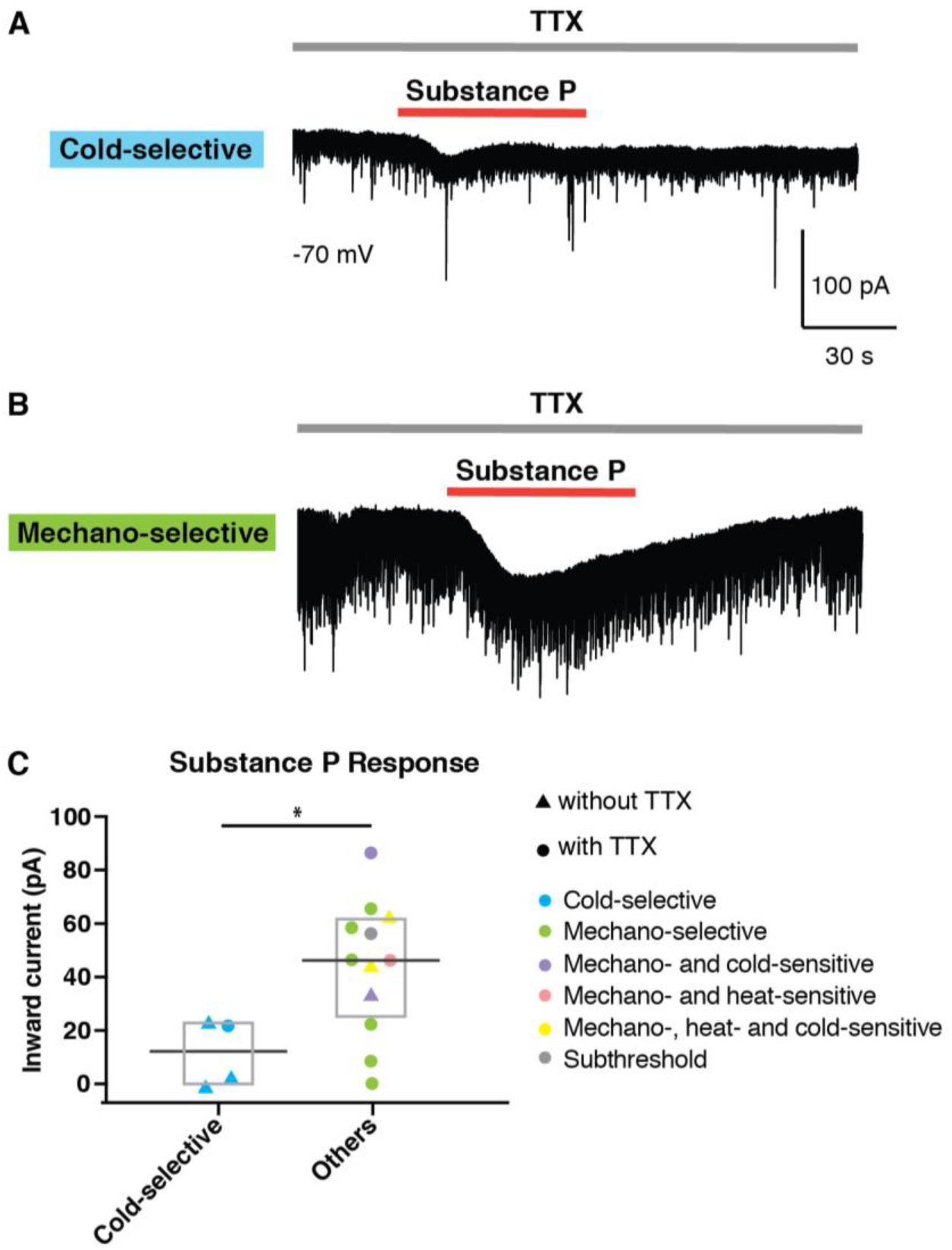
Cold-selective SPB neurons show little or no responsive to Substance P. **(A-B)** Representative traces of a cold-selective SPB neuron **(A)** and a mechano-selective SPB neuron **(B)** in response to bath application of Substance P (2µM) in the presence of tetrodotoxin (0.5 µM). **(C)** Amplitude of inward current of cold-selective and other SPB neurons induced by Substance P (2µM) in the presence or absence of TTX (0.5 µM), as indicated. Since cold-neurons showed little to no response to Substance P even in the absence of TTX, these data were pooled. Box plots are median and interquartile range with symbols representing data points from individual SPB neurons (n = 4 cold-selective and 12 other SPB neurons, * indicates p < 0.05, Wilcoxon test).

### Cold-selective SPB neurons have low membrane capacitance, high membrane resistance, and small soma size

To further investigate cold-selective SPB neurons, we examined their passive membrane properties. Cold-selective lamina I SPB neurons were not different than other SPB neurons with respect to resting membrane potential (Figure 4A). However, we found that cold-selective neurons had significantly lower membrane capacitance (Figure 4B) and higher membrane resistance (Figure 4C). These findings raised the possibility that cold-selective neurons are smaller in size than the others. To address this idea directly, we measured the soma area of SPB neurons that were filled with Alex 488 during recording. In agreement with capacitance and resistance measurements, the soma size of cold-selective SPB neurons was significantly smaller than that of other lamina I SPB neurons (Figures 4D and 4E).

**Figure 4.**
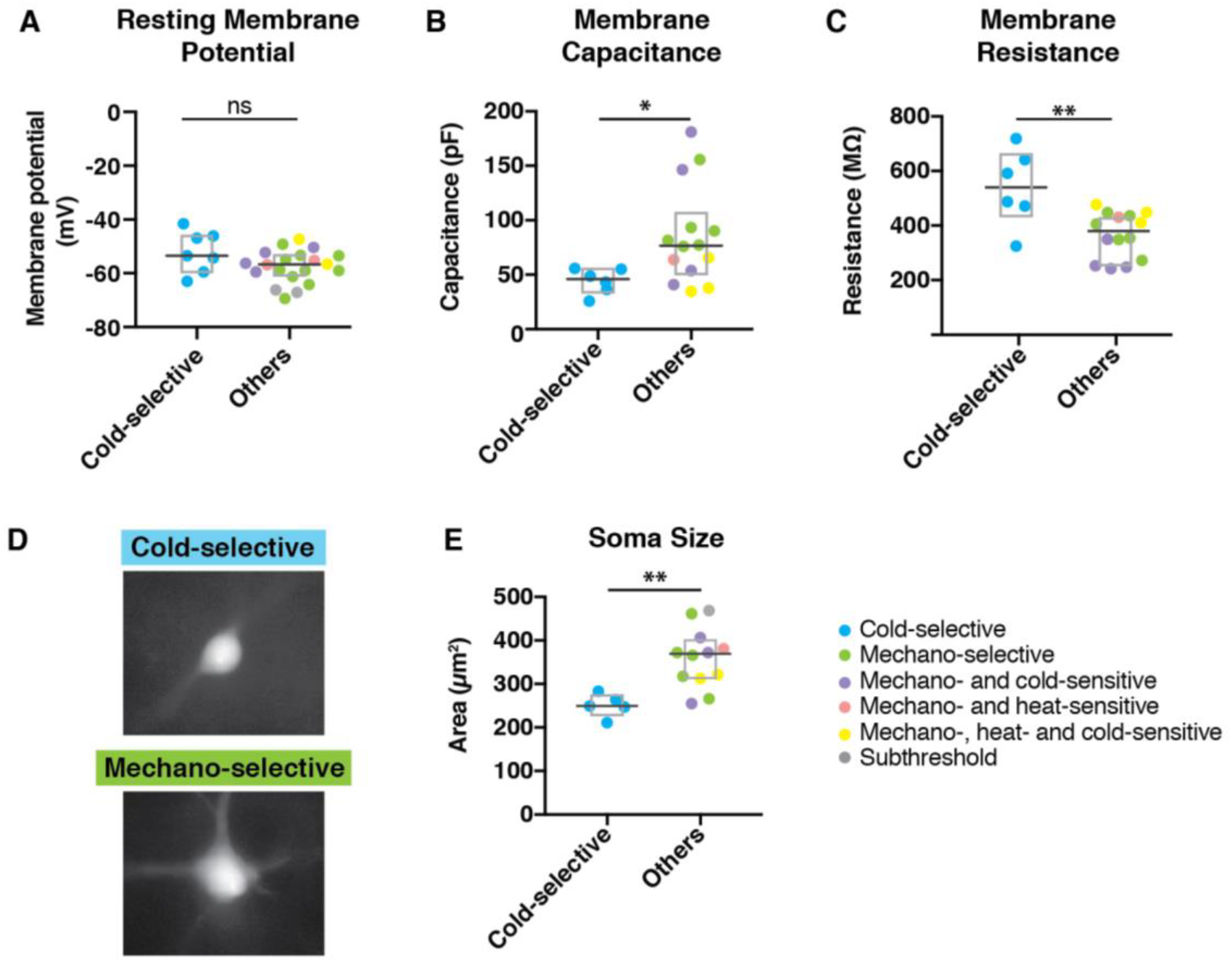
Cold-selective SPB neurons have distinct membrane properties and small soma size. **(A)** Resting membrane potential of cold-selective SPB neurons is not different than other SPB neurons. Box plots are median and interquartile range with symbols representing data points from individual SPB neurons (n = 7 cold-selective SPB neurons and 20 other SPB neurons; n.s., p > 0.05, Wilcoxon test). **(B)** Membrane capacitance of cold-selective SPB neurons is significantly lower than that of other SPB neurons. Box plots are median and interquartile range with symbols representing data points from individual SPB neurons (n = 6 cold-selective SPB neurons and 14 other SPB neurons; * indicates p < 0.05, Wilcoxon test). **(C)** Membrane capacitance of cold-selective SPB neurons is significantly higher than that of other SPB neurons. Box plots are median and interquartile range with symbols representing data points from individual SPB neurons (n = 6 cold-selective SPB neurons and 14 other SPB neurons; ** indicates p < 0.01, Wilcoxon test). **(D-E)** Representative images **(D)** and quantification **(E)** of soma size from cold-selective SPB neurons compared to other SPB neurons. Box plots are median and interquartile range with symbols representing data points from individual SPB neurons (n = 5 cold-selective SPB neurons and 12 other SPB neurons; ** indicates p < 0.01, Wilcoxon test).

### Cold-selective SPB neurons are distinct with respect to excitatory input

The finding that cold-selective SPB neurons have significantly higher membrane resistance than other SPB neurons raised the possibility that cold-selective neurons have fewer open channels. To address this idea in more detail, we quantified the number and size of spontaneous EPSCs (sEPSCs), recording in voltage clamp mode at a holding potential of −70 mV to isolate excitatory inputs. All cold-selective SPB neurons showed extremely sparse sEPSCs (Figures 5A and 5B), reflecting a very low level of basal input; in contrast, most of the non-cold selective SPB neurons received a continuous barrage of sEPSCs regardless of their functional response properties (Figures 5A and 5B). Thus, the median sEPSC frequency of cold-selective SPB neurons was significantly lower than that of non-cold neurons (4.1 vs 36.6 Hz), whereas the sEPSC amplitude was not different. These findings reinforce the idea that cold-selective SPB neurons are part of a distinct neural circuit that is quiet with respect to ongoing activity.

**Figure 5.**
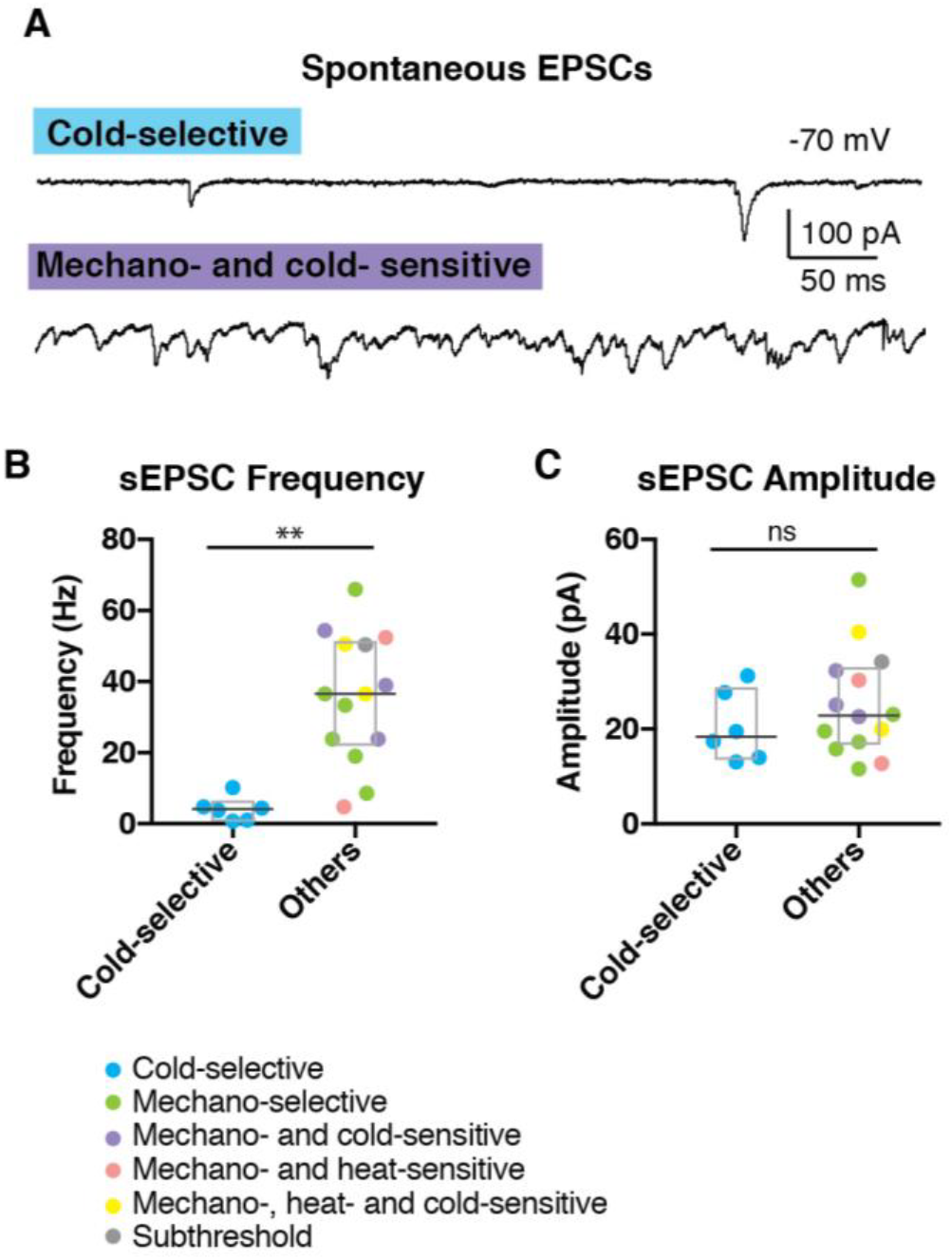
Cold-selective SPB neurons show few EPSCs. **(A)** Representative trace of cold-selective SPB neuron (upper trace) and mechano- and cold-sensitive SPB neuron (lower trace). (**B-C)** Quantification of sEPSC frequency **(B)** and amplitude **(C)** of SPB neurons. Box plots are median and interquartile range with symbols representing data points from individual SPB neurons (n = 6 cold-selective SPB neurons and 14 other SPB neurons; ** indicates p < 0.01, n.s. indicates p > 0.05, Wilcoxon test).

### Cold-selective SPB neurons belong to distinct spinal circuits

To further characterize this cold-selective circuit, we examined whether SPB neurons are distinct with respect to the types of inhibitory input they receive. Nos1 and Pdyn populations are two, largely non-overlapping subtypes of inhibitory neurons in the dorsal horn that are activated by noxious stimuli (Boyle et al., 2017; Iwagaki et al., 2013; Kardon et al., 2014; Polgár et al., 2013; Sardella et al., 2011). In behavioral experiments, Nos1 neurons have been implicated in the inhibition of heat and mechanical stimuli, whereas Dyn neurons have been implicated in the inhibition of itch and mechanical pain (Duan et al., 2014; Huang et al., 2018; Kardon et al., 2014; Ross et al., 2010). Whether either of these inhibitory neuron subtypes plays a role in the inhibition of cold is unknown.

To address this, we used double transgenic mice harboring Ai32, a Cre-dependent channel rhodopsin (ChR2), together with either Nos1-CreER or Pdyn-Cre alleles. First, we recorded from ChR2-expressing neurons confirm that optogenetic stimulation with blue light was sufficient to induce action potentials in Nos1 and Pdyn neurons (Figures 6A – 6C). Next, we recorded from SPB neurons in voltage clamp mode (holding potential = −40 mV) to record optogenically-induced currents that were observed upon activation of either Nos1-CreER or Pdyn-Cre populations (Figure 6D). Somewhat surprisingly, only one of 13 SPB neurons showed evidence of inhibitory input from Nos1-neurons, whereas 13 of 14 SPB neurons showed IPSCs upon stimulation of Pdyn-Cre neurons (Figure 6E). This finding suggested that, as general rule, SPB neurons receive direct inhibitory input from Pdyn neurons but not Nos1 neurons.

**Figure 6.**
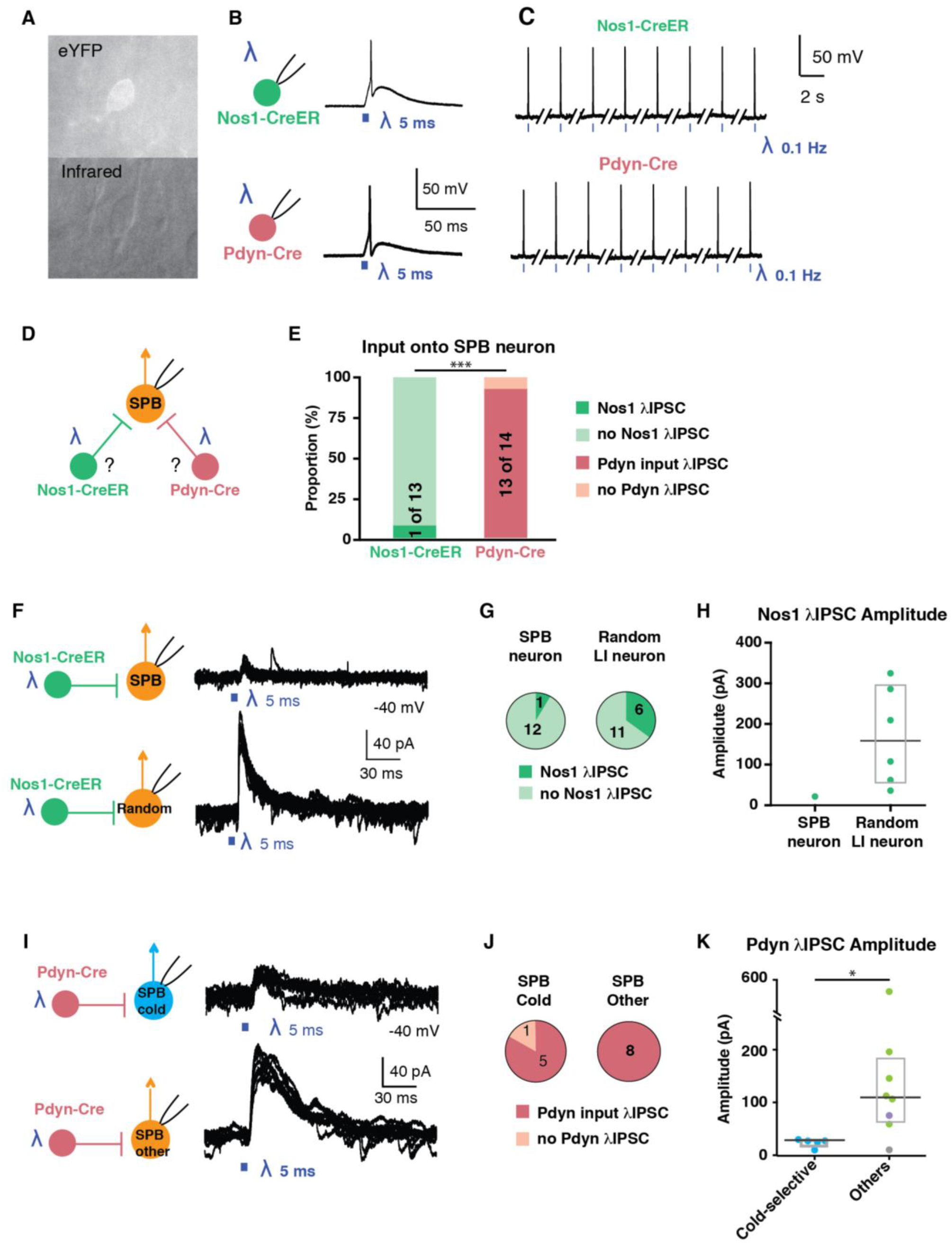
Cold-selective SPB neurons belong to distinct spinal circuits. **(A)** Fluorescent (top) and infrared (bottom) images of a genetically-labeled neuron that is targeted for patch-clamp recordings. **(B-C)** Optogenetic activation of Nos1-CreER (top) or Pdyn-Cre (bottom) neurons faithfully elicits action potentials upon single **(B)** or repeated stimulation at 0.1 Hz **(C)**. **(D)** Schematic illustrating experimental design to determine whether SPB neurons receive input from Nos1 or Pdyn inhibitory interneurons. **(E)** Proportion of SPB neurons that receive IPSCs in response to optogenetic activation of Nos1-CreER (green) or Pdyn-Cre inhibitory interneurons (red). *** indicates significantly different, p < 0.001, Fisher’s exact test. **(F)** Schematic and superimposed example traces of IPSCs of a SPB neuron (top) or a random neuron in lamina I (bottom) in response to optogenetic stimulation of Nos-CreER neurons. **(G)** Proportion of neurons that receive IPSCs in response to optogenetic activation of Nos1-CreER. **(H)** Amplitude of IPSCs observed upon optogenetic stimulation of Nos1-CreER neurons. Box plots are median and interquartile range **(I)** Schematic and superimposed example traces of IPSCs of a cold-selective SPB neuron (top) or other SPB neurons (bottom) in response to optogenetic stimulation of Pdyn-Cre neurons. **(J)** Proportion of SPB neurons that receive IPSCs in response to optogenetic activation of Pdyn-Cre neurons. **(K)** Amplitude of IPSCs observed upon optogenetic stimulation of Pdyn-Cre neurons. Box plots are median and interquartile range with symbols representing data points from individual SPB neurons (n = 5 cold-selective SPB neurons (blue), 6 mechano-selective neurons (green), 1 mechano- and cold-selective (purple) and 1 subthreshold (grey). * indicates significantly different, p < 0.05, Wilcoxon test.

To ensure that this apparent absence of synaptic input from Nos1 neurons onto SPB neurons was not simply a technical artifact, we next recorded random lamina I neurons (Figure 6F). We found that 6 of 15 random lamina I neurons showed IPSCs in response to optogenetic stimulation of Nos-CreER neurons (Figures 6F and 6G). Moreover, the IPSC amplitude in random neurons was much stronger (median = 158.7 pA, n = 6) than that observed in the singe SPB neuron that received input (20.8 pA, n=1). Thus, Nos1 neurons provide functional inhibitory input to some lamina I neurons, but not SPB neurons as a general class.

Next, we analyzed the Pdyn input in more detail (Figure 6I). We found that optogenetic stimulation of Pdyn neurons gave rise to large inhibitory post synaptic currents (IPSCs) in most SPB neurons, regardless of their response properties to natural stimulation of the skin (Figure 6E). However, although most cold-selective (5 of 6) and all non cold-selective (8 of 8) SPB neurons received IPSCs from Pdyn neurons, there was a significant difference with respect to the magnitude of this inhibition. The amplitude of optogenetically-induced IPSCs in cold-selective SPB neurons was significantly smaller than that observed in other SPB neurons (Figures 6E and 6F). These data suggest that, while Pdyn interneurons provide inhibitory input to many SPB neurons, their contribution to the inhibition of cold-selective SPB neurons is almost negligible.

## DISCUSSION

Our study provides evidence that cold is mediated by a subset of SPB neurons that have many distinctive properties including little to no response to Substance P, low capacitance, high resistance, small soma size, low basal drive, and a lack of inhibitory input from either Pdyn or Nos1 inhibitory neurons. Thus, these cells represent a distinct output channel through which cold information is conveyed from the periphery to the brain (Figure 7).

**Figure 7.**
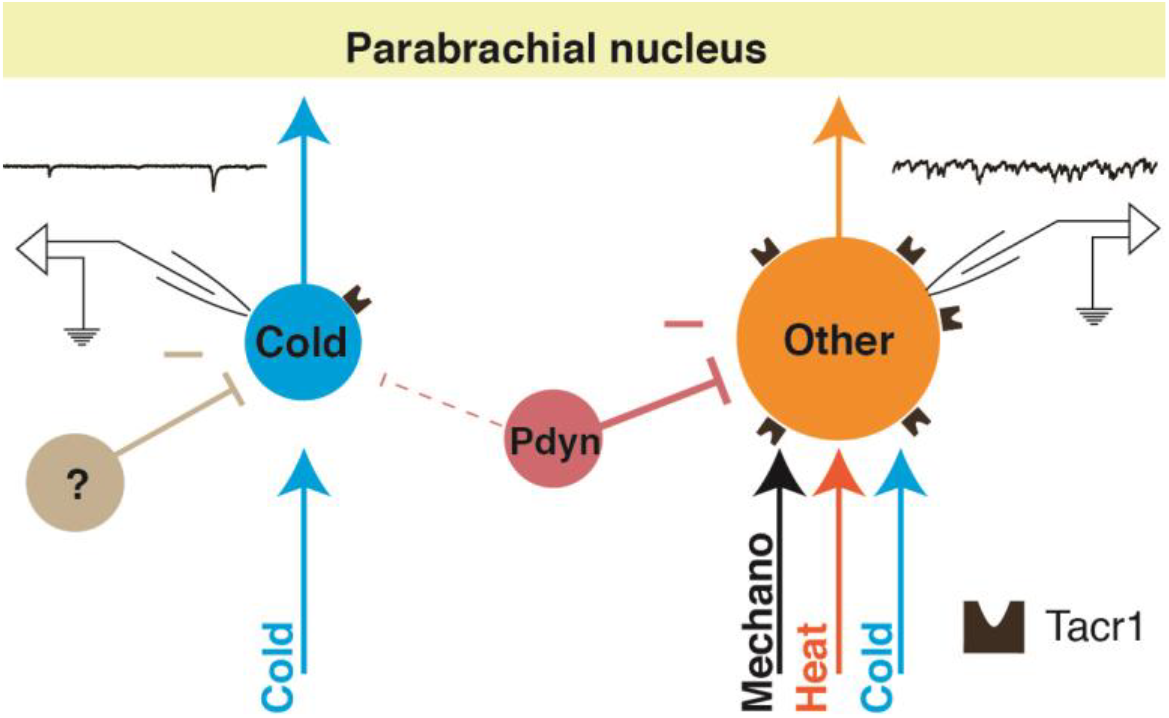
A labeled-line for cold from the periphery to the parabrachial nucleus. Cold-selective SPB neurons are distinct from other subtypes in multiple regards including selective input from cold afferents, smaller soma size, little to no expression of Tacr1, low basal drive, and little to no inhibition from Pdyn inhibitory interneurons.

The idea that cold is conveyed by a distinct output channel is not without precedent. Previous extracellular single unit recordings in cat, rat and monkey have shown that cold-selective STT neurons are distinct from polymodal nociceptive STT neurons with respect to thalamic projection patterns and conduction velocities (Craig et al., 2002, 2001; Craig and Dostrovsky, 2001; Craig and Hunsley, 1991; Craig and Kniffki, 1985; Dostrovsky and Craig, 1996; Zhang et al., 2006). Our study reveals that the SPB pathway, like the STT pathway, involves a specific channel for cold. Whether cold-selective SPB neurons and cold-selective STT neurons arise from distinct output neurons or from a common subset of spinal projection neurons with divergent collaterals remains to be addressed.

One of the intriguing findings of our study is that distinct spinal interneuron subtypes show differential connectivity. Specifically, our optogenetic experiments suggest that most SPB neurons do not receive direct inhibitory input from Nos1 neurons. However, there are likely exceptions to this rule since ‘giant cells’, which are an extremely sparse population of Tacr1-negative SPB neurons of unknown function, have a very large number of synaptic puncta from Nos1 neurons (Puskár et al., 2001). A second population of inhibitory neurons, Pdyn cells, provides inhibitory input to the majority of SPB neurons in a manner that shows strong modality specificity: some SPB neurons receive strong input, but very little is observed in those that are cold-selective. Since cold-selective SPB neurons have spontaneous IPSCs (data not shown) but receive little to no input from either Nos1 or Pdyn interneurons, this raises the question as to which inhibitory interneuron subtype(s) mediate the inhibition of cold.

The results from our study are in good agreement with previous studies and lead to several new predictions. For instance, our study reveals that cold-selective SPB neurons have a relatively small soma size and show fewer EPSCs compared to other subtypes. It has been reported that small SPB neurons have fewer excitatory synapses and express the Gria1 subunit of the AMPA receptor (also known as GluA1), whereas large SPB neurons have numerous excitatory synapses and express Gria4 (also known as GluA4) (Polgár et al., 2010). These observations raise the interesting possibility that cold-selective SPB neurons express Gria1, rather than Gria4, thereby enabling distinct types of synaptic plasticity. Previous studies have also suggested that cold-selective (albeit random) lamina I neurons in cat are pyramidal (Han et al., 1998), and lamina I pyramidal neurons have been shown to be negative for Tacr1-immunoreactivity in rat and monkey (Almarestani et al., 2007; Saeed et al., 2015; Saeed and Ribeiro-da-Silva, 2013; Yu et al., 2005, 1999). Thus, our finding that cold-selective SPB neurons show little to no response to Substance P is consistent with the possibility that they may be pyramidal in shape. In future studies, it will be important to perform detailed morphological analyses and comprehensive neurochemical profiling to tests these predictions.

We and others have recently shown that cutaneous Trpm8 sensory neurons are cold-responsive neurons with several unique characteristics (Dhaka et al., 2007; Jankowski et al., 2017; Knowlton et al., 2013). For example, as a group they have the fastest conduction velocities of all cutaneous C-fibers, are capable of firing at very high frequencies, and have cell bodies among the smallest in the DRG (Jankowski et al., 2017). However, it should be noted that Trpm8 neurons likely represent more than one subtype of afferent, since ¾ are unimodal (responding to cold alone), whereas ¼ are polymodal (responding to mechanical and cold stimuli) (Jankowski et al., 2017). Our data suggest that unimodal Trpm8 afferents are likely those that provide input onto cold-selective SPB neurons. This cold-only pathway, comprising a subset of Trmp8 cells together with cold-selective SPB neurons, forms the cellular basis for true labeled-line for cold from the periphery to the brain.

Presumably, such a specialized circuit for cold provides a distinctive sensory function. Cold-selective afferents have low basal activity and are exquisitely sensitive to rapid changes in temperature (Jankowski et al., 2017). Thus, high frequency activity within this circuit likely signals cooling. As an event detector, such activity would alert the organism of a shift in its environment, which might be pleasant or aversive, depending on the circumstance. Thereafter, the ongoing activity observed in response to a prolonged cold stimulus might provide temperature information that is critical for thermoregulation. Indeed, SPB neurons are known to regulate body temperature through downstream connections to preoptic area of the hypothalamus (Morrison and Nakamura, 2018). Thus, in a cold environment, activity in cold-selective SPN neurons could provide the sensory input that elicits heat production as well as the behavioral drive to seek warmth (Hammel and Pierce, 2003; Nagashima et al., 2000; Romanovsky, 2006).

## METHODS

### Animals

Five-to eight-week-old mice of both sexes were used in this study. Wild type mice (C57BL/6) were purchased from Charles River (Horsham, PA). Genetically modified mice were purchased from The Jackson Laboratory (Bar Harbor ME) and bred in house. These were: Pdyn-IRES-Cre, a non-disruptive Cre recombinase knock-in at the endogenous prodynorphin locus (stock: 027958), Nos1-CreER, a disruptive Cre recombinase knockin to the endogenous neuronal nitric oxide synthase locus (stock: 014541), and Ai32, a Cre-dependent ChR2 fusion protein, ChR2(H134)/EYFP inserted into the Rosa locus (stock: 012569). Tamoxifen (Sigma, 0.4 mg/kg; IP) was injected into mice harboring the Nos1-CreER allele at post-natal day 14 (P14), three weeks prior to electrophysiological experiments. Mice were given free access to food and water and housed under standard laboratory conditions. The use of animals was approved by the Institutional Animal Care and Use Committee of the University of Pittsburgh.

### Stereotaxic injection of DiI

Four-to six-week-old mice were anesthetized with isoflurane and placed in a stereotaxic apparatus. A small hole was made in the skull bone with a dental drill. A glass pipette was used to inject 100 nl of FAST DiI oil (2.5 mg/ml; Invitrogen, Carlsbad, CA) into the left lateral parabrachial area (relative to lambda: anteroposterior −0.5 mm; lateral 1.3 mm; dorsoventral −2.4 mm). The head wound was closed with stitches. After recovery from the anesthesia, the animals fed and drank normally. DiI was injected at least five days prior to electrophysiological recordings.

### Semi-intact somatosensory preparation

Semi-intact somatosensory preparation was made as previously described with small modification (Hachisuka et al., 2016). Briefly, young adult mice (5–9 weeks old) were deeply anesthetized and perfused transcardially through the left ventricle with oxygenated (95% O2 and 5% CO2) sucrose-based artificial cerebrospinal fluid (ACSF) (in mM; 234 sucrose, 2.5 KCl, 0.5 CaCl2, 10 MgSO4, 1.25 NaH2PO4, 26 NaHCO3, 11 Glucose) at room temperature. Immediately following perfusion, the skin was incised along the dorsal midline and the spinal cord was quickly exposed via dorsal laminectomy. The right hindlimb and spinal cord (~C2 – S6) were excised, transferred into Sylgard-lined dissection/recording dish, and submerged in the same sucrose-based ACSF, which circulated at 50 ml/min to facilitate superfusion of the cord. Next, the skin innervated by the saphenous nerve and the femoral cutaneous nerve was dissected free of surrounding tissue. L2 and L3 DRG were left on the spine. Dural and pial membranes were carefully removed and spinal cord was pinned onto the Sylgard chamber with the right dorsal horn facing upward. Following dissection, the chamber was transferred to the rig. Then the preparation was perfused with normal ACSF solution (in mM; 117 NaCl, 3.6 KCl, 2.5 CaCl2, 1.2 MgCl2, 1.2 NaH2PO4, 25 NaHCO3, 11 glucose) saturated with 95% O2 and 5% CO2 at 31 °C. Tissue was rinsed with ACSF for at least 30 min to wash out sucrose. Thereafter, recordings were performed for up to 6 h post-dissection.

### Patch clamp recording from dorsal horn neurons

Neurons were visualized using a fixed stage upright microscope (BX51WI Olympus microscope, Tokyo, Japan) equipped with a 40× water immersion objective, a CCD camera (ORCA-ER Hamamatsu Photonics, Hamamatsu City, Japan) and monitor screen. A narrow beam infrared LED (L850D-06 Marubeni, Tokyo, Japan, emission peak, 850 nm) was positioned outside the solution meniscus, as previously described (Hachisuka et al., 2016; Safronov et al., 2007; Szucs et al., 2009). Projection neurons in lamina I were identified by DiI fluorescence following injection into the lateral parabrachial nucleus. Whole-cell patch-clamp recordings were made with a pipette constructed from thin-walled single-filamented borosilicate glass using a microelectrode puller (PC-10; Narishige International, East Meadow NY). Pipette resistances ranged from 6 to 12 MΩ. Electrodes were filled with an intracellular solution containing the following (in mM): 135 K-gluconate, 5 KCl, 0.5 CaCl2, 5 EGTA, 5 HEPES, 5 MgATP, pH 7.2. Alexa fluor 488 (Invitrogen; 25 µM) was added to confirm recording from the target cell. Signals were acquired with an amplifier (Axopatch 200B, Molecular Devices, Sunnyvale CA). The data were low-pass filtered at 2 kHz and digitized at 10 kHz with an A/D converter (Digidata 1322A, Molecular Devices) and stored using a data acquisition program (Clampex version 10, Molecular Devices). The liquid junction potential was not corrected.

### Natural stimulation to the skin

To search for a cell’s receptive field, a firm brush or a 4 g von Frey filament was applied systematically over the skin. If no response to mechanical stimulation was observed, then hot (50 °C) or cold (0 °C) saline was applied in pseudorandom order across the skin. Once a receptive field was located, stimuli were reapplied directly to the receptive field for 1 s. For mechanosensitive neurons, a variety of mechanical stimuli were applied (small firm paintbrush and/or von Frey filaments (1, 2 and 4 g), but these data were pooled for in this study for simplicity. Thermal stimulation was applied using 1 ml of hot (50 °C) or cold (0 °C) saline applied gently to the receptive field over 1 s using 10 cc syringe and 18 G needle. For current clamp recordings, the action potential frequency was calculated in 1 s bins. Responses to natural stimuli were considered significant if the number of action potentials during the stimulation period (1 s) were more than 3 times the standard deviation of the baseline period averaged over preceding 9 s. For voltage clamp recordings, EPSCs were detected by MiniAnalysis (Synaptosoft). EPSC frequency for during the stimulation (1 s) was calculated. Baseline values were averaged over 9 s before stimulation.

### Quantification of soma size

For quantification of soma size, Alexa 488 filled neurons were imaged using Micro-manager (an open source software program) with a 40× objective. The area of the soma was analyzed in ImageJ, tracing the edge of the soma.

### Optogenetic stimulation

For optogenetic stimulation, a blue light pulse (GFP filter, centered around 485 nm, Lambda DG-4, Sutter instruments) was applied through the objective (40×) of the microscope for 5 ms using a shutter that was controlled by Clampex software (Clampex version 10, Molecular Devices). The light power on the sample was 1.3 mWmm^−2^. The peak amplitude of outward current induced by blue light stimulation was measured in voltage clamp mode at −40 mV.

### Pharmacology

The drugs (tetrodotoxin: Tocris, Substance P acetate salt hydrate: Sigma) were dissolved in ACSF and applied by exchanging solutions via a three-way stopcock using a modified chamber adapted for pharmacological experiments that limits drug dilution (Hachisuka et al., 2016).

## ACKNOWLEDGMENTS

We thank Michael S. Gold for helpful comments. Research reported in this publication was supported by the National Institute of Arthritis and Musculoskeletal and Skin Diseases of the National Institutes of Health under Award Number R01AR063772 to S.E. Ross) and the National Institute of Neurological Disorder and Stroke of the National Institutes of Health under Award Number R01 NS096705 to H.R. Koerber.

